# A high-throughput fluorescence-based assay for rapid identification of petroleum degrading bacteria

**DOI:** 10.1101/498345

**Authors:** K. E. French, N. Terry

## Abstract

Over the past 100 years, oil spills and long-term waste deposition from oil refineries have significantly polluted the environment. These contaminants have widespread negative effects on human health and ecosystem functioning. Natural attenuation of long chain and polyaromatic hydrocarbons is slow and often incomplete. Bioaugmentation of polluted soils with indigenous bacteria that naturally consume petroleum hydrocarbons could speed up this process. However, the characterization of bacterial crude oil degradation efficiency—which often relies upon expensive, highly specialized gas-chromatography mass spectrometry analyses--can present a substantial bottleneck in developing and implementing these bioremediation strategies. Here, we develop a low-cost, rapid, high-throughput fluorescence-based assay for identifying wild-type bacteria that degrade crude oil using the dye Nile Red. We show that Nile Red fluoresces when in contact with crude oil and developed a robust linear model to calculate crude oil content in liquid cell cultures based on fluorescence intensity (FI). To test whether this assay could identify bacteria with enhanced metabolic capacities to break down crude oil, we screened bacteria isolated from a former Shell Oil refinery in Bay Point, CA and identified one strain (*Cupriavidus* sp. OPK) with superior crude oil depletion efficiencies (up to 83%) in only three days. We further illustrate that this assay can be combined with fluorescence microscopy to study how bacteria interact with crude oil and the strategies they use to degrade this complex substance. We show for the first time that bacteria use three key strategies for degrading crude oil: biofilm formation, direct adherence to oil droplets, and vesicle encapsulation of oil. We propose that the quantitative and qualitative data from this assay can be used to develop new bioremediation strategies based on bioaugmentation and/or biomimetic materials that imitate the natural ability of bacteria to degrade crude oil.

## Introduction

Over 50 million tons of crude oil have been spilled into the ocean and on land as a result of equipment failure, transportation accidents, and human tampering since the 1970s (Jernelöv 2010; ITOPF 2017). Such incidents negatively impact local ecosystems and decrease biodiversity in contaminated areas (Azevedo-Santos 2016; Das and Chandran 2011; Turner and Renegar 2017). As many of these compounds are carcinogens (e.g. pyrene, benzene), oil spills also present an unprecedented threat to human health. High molecular weight polyaromatic hydrocarbons (PAHs) like benzo(a)anthracene, benzo(a)pyrene, and benzo(g,h,i)perylene can accumulate to toxic levels in the air (de Gouw et al. 2011; Srogi 2007). Petroleum hydrocarbons can also contaminate land and water reservoirs, making these areas unsuitable for agriculture or settlement (Maliszewska-Kordybach dand Smreczak 2000). Particularly in developing countries, these risks go unmitigated. For example, after a series of spills from Shell Oil pipes and refineries in the Niger Delta from the 1990s to early 2000s, drinking wells are currently contaminated with benzene levels that are 900x the safe level (Lindén and Pålsson 2013).

Although the number of spills has declined over the past ten years, clean-up of historic spills is non-existent or often incomplete (Jernelöv 2010). Ocean spills rely on burning or mopping-up crude oil and relying on native ocean bacteria to break down the remaining petroleum hydrocarbons (Kostka et al. 2011). Shoreline and inland spills are more difficult to clean up effectively. Environmental hazard teams often rely on chemical washing or removal of contaminated sand or soil, and where this is not possible, natural attenuation of petroleum hydrocarbons by native organisms (Das and Chandran 2011; Li et al. 2016). However, natural processes of degradation are often slow and only remove shorter-chain hydrocarbons. Long chain hydrocarbons and PAH often resist degradation because they are inaccessible to organisms (due to mineralization, hydrophobicity, toxicity, and adsorption onto soil particles) and only a small number of bacteria have the metabolic capacity to degrade these compounds (Bamforth and Singleton 2005).

Bioremediation using bacteria isolated from polluted sites may enhance the speed and efficiency of petroleum hydrocarbon removal (Bento et al. 2005; Das and Chandron 2011). The advantage of using indigenous bacteria is two-fold: these bacteria are already equipped to handle local environmental conditions and it gets around regulatory hurdles which may prohibit the introduction of foreign (or genetically engineered) bacteria into the area (Thompson et al. 2005; Urgun-Demirtas et al. 2006). Known bacterial petroleum hydrocarbon degraders include *Alcanivorax borkumensis*, *Bacillus subtilis*, *Burkholderia cepacia*, *Pseudomonas fluorescens*, *Pseudomonas marginalis*, and *Pseudomonas oleovorans* (Brooijmans et al. 2009; Rojo et al. 2009). These bacteria contain monooxygenases and dioxygenases which insert oxygen into petroleum hydrocarbons to break down their carbon structure (Wang and Shao 2013). Some bacteria also produce secondary compounds, namely biosurfactants, which make crude oil more accessible (Mohanty et al. 2013). Biosurfactants reduce the surface tension between the oil and water interface and allow recalcitrant compounds, like PAHs, to precipitate into the aqueous phase, where they can then be metabolized by bacteria.

Inocula can be made of selected petroleum degrading bacteria and then applied to polluted land as a form of bioaugmentation. However, screening, identification, and characterization can be a time-consuming and expensive process. Traditional methods to characterize bacterial degradation of crude oil rely on gas chromatography mass spectrometry (GC/MS). This method requires specialized compound databases and the analytical capacity to analyze crude oil-which many mass spectrometry labs do not have. Few commercial companies will analyze crude oil from cell culture, and when this is possible, large volumes of culture are needed and tests can cost >$400 per sample for custom analyses. From any given site, over 100 strains of bacteria might be isolated. The cost of such analyses could be prohibitive in cases where funding is limited. For example, analysis of 100 strains with three replicates at $470 per sample would be $141,000.

To speed up the process of discovery, we sought to exploit the optical properties of the dye Nile Red to develop a high-throughput fluorescence-based assay that uses 96-well microtiter plates and a plate reader to detect bacteria that can degrade petroleum hydrocarbons. Nile Red is a colorless compound which fluoresces red when in contact with hydrophobic substances (Greenspan and Fowler 1985). It is often used as a stain for lipids and fatty acids (Greenspan et al. 1985; Rumin et al. 2015; Shrivastav et al. 2010), but has never been used, to our knowledge, as a way to measure the amount of crude oil in cell culture media or in environmental samples.

Our research had three key objectives: (1) to determine whether there was a correlation between fluorescence intensity (FI) and crude oil content, (2) to develop a new model to calculate crude oil content based on FI, and (3) to identify whether this assay could detect bacterial strains isolated from the natural environment that are able to degrade crude oil. Based upon statistical analysis of assay data, we show that this assay can measure and calculate crude oil degradation by wild-type bacteria down to the level of 1.25 nl/μl in as little as three days. We also suggest that this assay can be a powerful new tool to study *how* bacteria degrade crude oil. When combined with fluorescence microscopy, we show that bacterial mechanisms for dispersing, sequestering, and degrading crude oil can be observed and analyzed. We anticipate that this assay can be used not only to rapidly detect novel bacteria for bioremediation, but to also advance the development of new bio-inspired solutions to remove petroleum hydrocarbons from the environment.

## Results & Discussion

To determine whether the fluorescence intensity of Nile-Red stained crude oil was correlated with crude oil content, we set up 96-well plates with two serial dilutions of crude oil (1 μl - 16 μl, and 250 nl – 4 μl, equivalent to 1-8% and 0.125%-2% crude oil respectively) as described in the method section. Each plate received four treatments: LB without bacteria, LB with bacteria, MSM without bacteria and MSM with bacteria. For these initial proof-of-concept assays, we randomly selected a bacterial strain (*Cupriavidus* sp. strain OPK) from our freezer stocks as a model bacterium. At day 0 (T_0_), there was a strong correlation between fluorescence intensity (FI) and optical density (OD) for all treatments (cells vs. no-cells with LB or MSM) (Fig. 1A-B). This indicates FI is a strong proxy for crude oil content. After T_0_, for the wells containing bacteria the correlation between FI and OD varied as cells proliferated and the amount of crude oil declined.

**Figure 1:**
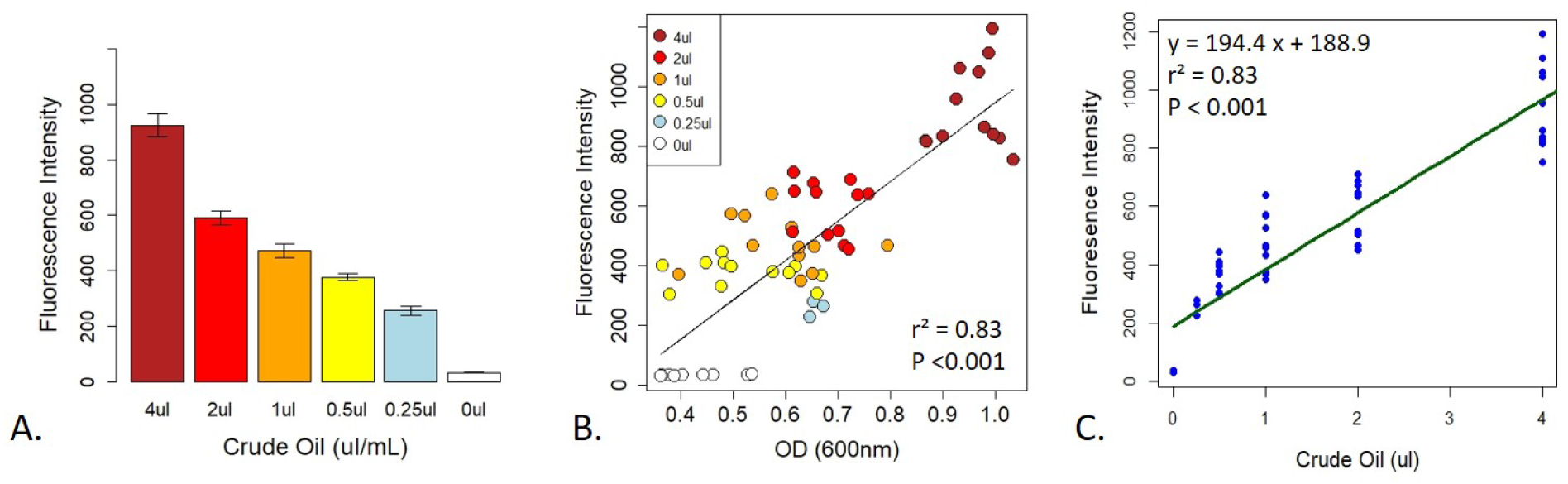
Correlation between fluorescence intensity and crude oil content. There was a strong relationship between fluorescence intensity and crude oil content when crude oil was incubated with Nile Red (A). This relationship was confirmed by the strong correlation between crude oil fluorescence intensity and optical density on D_0_ (B). Figure 1.C represents the linear model calibrating fluorescence intensity with crude oil content. All fluorescence intensity data is based on excitation at 535nm and emission at 650nm.

To determine the strength of the relationship between FI and crude oil content, we created several linear models from the T_0_ data for each of the treatments (described fully in the method section). The final model used for calibration was based on the MSM with bacteria treatment (Fig. 1C). The model had an R^2^ of 0.83 with a y-intercept of 188.9 ± 22.37 and a concentration coefficient of 194.4 ± 11.11 with a residual standard error of 124 on 61 degrees of freedom (F_1,61_= 305.9, p <0.0001). The standard errors and their fitted values were randomly distributed, the residual errors were normally distributed in the Q-Q plot, and Cooks’ Distances among all points was less than 1 (**SI Fig. 1**). The final calibration equation is y = 194.4x + 188.9. To calculate crude oil depletion efficiency, we developed an additional approach based on the following equation: DE = FIT_0_-FIT_3_/FIT_0_, where DE is the ‘depletion efficiency’ and FI_x_ is the fluorescence intensity on day x. The calibration curve and depletion efficiency equation thus provide two approaches to detect and quantify bacterial crude oil degradation.

To optimize assay parameters, we conducted a series of experiments to determine whether cell culture media or the presence of bacteria could alter the fluorescence of Nile Red-stained crude oil, leading to an over or under estimation of crude oil content. We found media type did not interfere, in terms of fluorescence quenching or overlapping in excitation/emission spectra, with the fluorescence of the Nile-Red stained crude oil. T-tests showed that there was no difference in fluorescence intensity in wells filled with bacteria and LB or MSM (*t* = 1.27, df = 30.25, p = 0.21) or in wells without bacteria in LB or MSM (*t* = 0.88, df = 33.26, p = 0.38). This indicates that cell and media autofluorescence is minimal and does not over-lap with the crude oil fluorescence. Although media-type did not affect the fluorescence of the Nile Red, we found that using Minimal Salt Media in the assays may provide a more accurate measure of bacterial crude oil degradation. The microbial rate of crude oil degradation was almost 10% greater in the presence of MSM. *Cupriavidus* sp. strain OPK grown in LB degraded on average 74% of the crude oil while *Cupriavidus* sp. strain OPK in MSM reduced crude oil by 83% (*t* = −2.8, df = 3.96, p = 0.04); however, there was no difference in the final biomass between the two treatments (*t* = −1.78, df = 2.77, p = 0.18). (**SI Fig. 2**). This is likely due to the fact that in MSM crude oil is the only source of nutrients while in LB there are other nutrients (e.g. amino acids from yeast extract) which could support bacterial metabolism.

To further test whether this assay could be used to distinguish differences in the efficiency of bacterial degradation of crude oil, we tested three strains of bacteria previously isolated from Shell Pond, a petroleum contaminated site in Bay Point, CA (Fig. 2). The site was a former Shell Refinery, where petroleum byproducts and other chemicals were deposited from the 1950s until the 1970s. Of the three bacterial species, two species (*Cupriavidus* sp. strain OPK and *Rhodococcus erythropolis* strain OSPS1) performed the best, depleting 69 ± 0.03% and 62 ± 0.03% of crude oil, respectively. The *Pseudomonas* sp. strain BSS only depleted 49 ± 0.03% of the crude oil. In contrast, minimal degradation was seen in control experiments using *E.coli* DH5α (loss of crude oil was between 1-3%, the same amount seen in the control wells with only crude oil and no bacteria). Although we used this assay to quantify bacterial efficiency at degrading crude oil under neutral circumstances, it could also be used in the future to look at how bacteria degrade crude oil under a gradient of different conditions (such as pH, temperature, or biostimulants) or other ecological variables like competition.

**Fig. 2.**
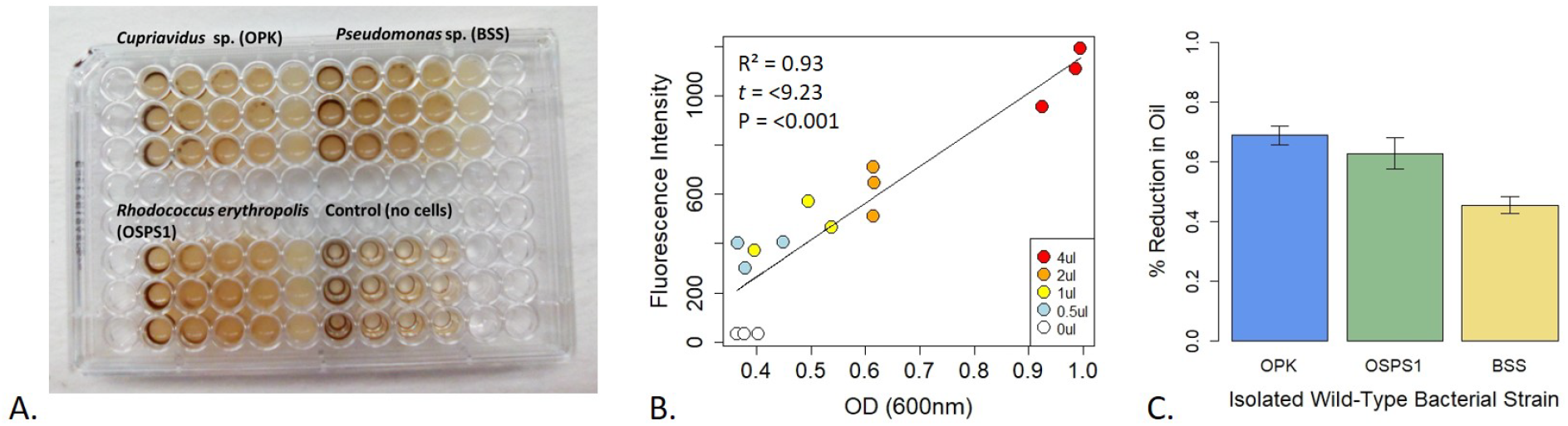
Design and performance of a fluorescence-based assay to detect bacteria that degrade crude oil. A. Assay of three bacteria isolated from Shell Pond after 3 days of exposure to crude oil (concentrations ranging from 0.125% - 2%) in Minimal Salt Media (MSM). Consumption of crude oil can be seen in wells with bacteria. B. Correlation between fluorescence intensity (FI) and optical density (OD). Each point in the plot represents the treatments from 2A (decreasing from 4 μl to 0.5 μl per well, for a concentration of 2 to 0.125 % crude oil per 200 μl well volume) at T_0_. Correlation between FI and OD for each treatment was >0.9 at T_0_, demonstrating the replicability of this assay. C. Amount of crude oil consumed by each bacterial strain when grown in 2% crude oil.

We also found that this assay lends itself well to investigating how bacteria interact with crude oil. For the above experiments, we extracted 1 μl of each bacterial strain from the 96-well plate at T_3_, T_10_, and T_20_ for fluorescence microscopy analysis. We found that each bacterial strain used a different strategy to sequester and break down crude oil.

The two most efficient strains relied on direct metabolism of crude oil. *Cupriavidus* sp. strain OPK formed biofilm network for long-distance transport of crude oil (Fig. 3A). These biofilms attached to the crude oil floating on the surface of the cell culture media and were anchored to the bottom of the microtiter well. Transport of crude oil through a network may allow for the efficient diffusion of petroleum hydrocarbons across highly specialized membranes embedded with monooxygenases and dioxygenases that break down petroleum hydrocarbons, such as alkB and p450cam (Gkorezis et al. 2016; French et al. *in prep*). Petroleum compounds degraded in this manner would be dispersed within the biofilm community. This efficient mode of crude oil dispersion and consumption may explain why this strain was the most effective at degrading crude oil. In contrast, *Rhodococcus erythropolis* strain OSPS1 attached directly to crude oil droplets (Fig. 3B). Other studies have shown that bacteria, such as *Alcanivorax borkumensi*, can attach to crude oil using exopolysaccharides and pili (Broojimans et al. 2009). This approach to degradation benefits individual bacteria: the compounds they metabolize go directly to their own growth and development. Crude oil is a complex substrate made of thousands of compounds, some of which may be toxic to bacteria (Xu et al. 2018). Potentially, both strains are able to withstand exposure to toxic elements while selectively metabolizing certain compounds.

**Figure 3.**
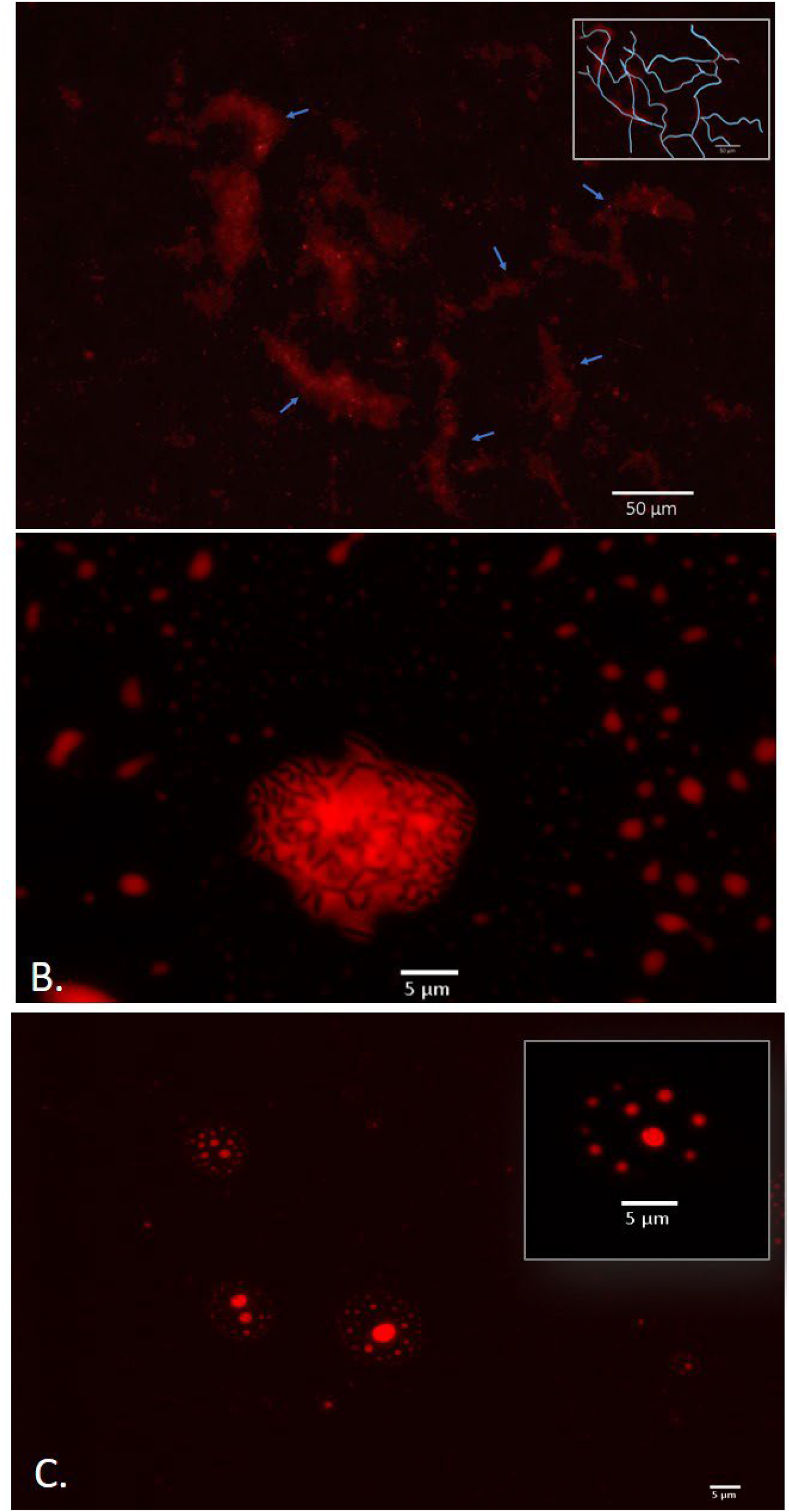
Bacterial strategies for crude oil degradation. A. *Cupriavidus* sp. OPK transported crude oil through biofilm networks after exposure to 1-12.5% crude oil for 3 days. Blue arrows point to channels within the network; the inset box shows a rough tracing of the network for clarity. B. *Rhodococcus erythropolis* strain OSPS1 (black rods) adhered to crude oil droplets. The oil droplet depicted is 21.03 um in diameter. C. *Pseudomonas* sp. strain BSS produced biosurfactants when exposed to crude oil which spontaneously formed vesicles. The vesicle depicted inset is 11.31 μm in diameter and contains small oil droplets ranging in size from 790 nm - 1.33 μm in diameter. All images are false-colored red; detailed description of image acquisition can be found in the Methods section.

In contrast, *Pseudomonas* sp. strain BSS relied on external metabolism of crude oil by encapsulating crude oil into vesicles. These vesicles were 4.17 −12.34 μm in diameter which contained small oil droplets ranging from 0.11 to 2.98 μm in diameter (Fig. 3C; **SI Fig. 3A**). In a few cases, these vesicles also contained bacteria (**SI Fig. 3B**). We believe these vesicles spontaneously form when *Pseudomonas* sp. strain BSS releases biosurfactants. We have observed a similar phenomenon in other wild-type strains of bacteria that produce biosurfactants (**SI Fig. 4**) and we know that *Pseudomonas* species produce rhamnolipids during the degradation of petroleum hydrocarbons (Hua and Wang 2014; Mulligan and Gibbs 2004). Our follow-up experimental research also confirms that biosurfactants can spontaneously form vesicles 5-150 μm in diameter in the presence of water and crude oil (**SI Fig. 5**). Surfactant-based degradation of crude oil offers several benefits: crude oil is broken up into smaller, easier to degrade droplets; PAHs may become more soluble in water; and a barrier is placed between the bacteria and potentially harmful compounds in crude oil.

Apart from advancing our fundamental knowledge of how bacteria degrade crude oil, the qualitative data garnered from this assay could be used in conjunction with the quantitative data on bacterial crude oil depletion efficiencies to select specific strains for different applications. For example, strains like *Cupriavidus* sp. strain OPK and *Rhodococcus erythropolis* strain OSPS1 might be more suitable for direct application to polluted soils as inoculum. They could also be applied in large quantities to bioreactors containing polluted soil (Robles-González et al. 2008). In contrast, strains like *Pseudomonas* sp. BSS might be more suitable for clean up of soils and water dominated by recalcitrant PAHs, where production of biosurfactants would be an asset (Bezza and Chirwa 2016).

This qualitative data could also be used to develop *de novo* bio-inspired solutions for bioremediation. Further genetic, molecular, and ecological analysis of bacteria screened using the Nile Red assay could lead to new ways to augment these capabilities or to develop new biomaterials that can degrade target toxins in the environment (Fig. 4). For example, research on the relationship between biofilm formation and petroleum degradation could lead to bioinspired filters which could be used for water purification. Similarly, understanding the structure, chemical composition and function of biosurfactant-based vesicles could be used to create synthetic vesicles for enhanced removal of recalcitrant PAHs from highly polluted environments where living inoculum might not survive. Increased academic and industrial interest in using synthetic biology to develop bio-based solutions to global challenges like environmental pollution mean that such solutions are on the horizon (French 2019).

**Figure 4:**
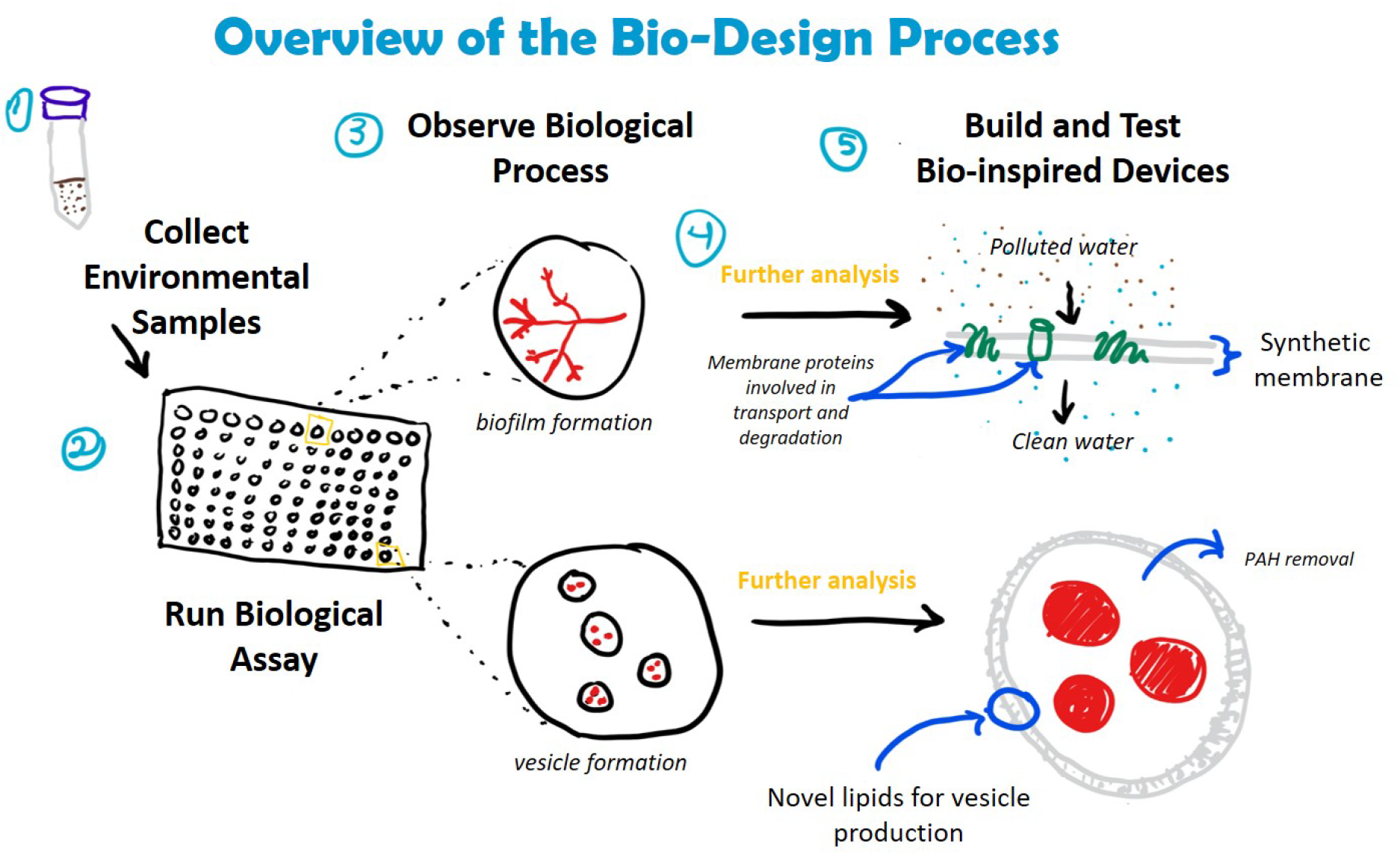
Incorporation of crude oil degradation assay into bio-design processes. Understanding how bacteria degrade crude oil could lead to the development of novel materials and devices for bioremediation. Examples include the creation of biofilters for water filtration (embedded with enzymes involved in hydrocarbon transport and degradation) and synthetic vesicles composed of bacterial-derived lipids for the removal of PAHs from complex hydrocarbon mixtures. The assay is compatible with a number of downstream analyses, including DNA extraction (e.g. to identify bacterial species, genes involved in hydrocarbon degradation) and mass spectrometry (e.g. LC/MS and MALDI to identify membrane proteins and lipids).

## Conclusion

Toxic compounds released during crude oil spills and waste from crude oil refineries threaten ecosystem functioning, local biodiversity, and human health. The versatile assay we have developed here will hasten our ability to identify bacteria that efficiently degrade petroleum hydrocarbons and could lead to the development of new bio-inspired solutions to cleaning up oil spills. Future research could develop variations on this fluorescence-based assay to rapidly identify bacteria that can degrade single hydrocarbon substrates (e.g. pyrene) or other classes of toxic environmental compounds (e.g. polychlorinated biphenyls, PCBs). Consequently, we anticipate that the assay system presented here could be used and modified by microbiologists, ecologists and engineers for a variety of applications within the fundamental and applied sciences.

## Methods

### Growth of bacterial strains

Bacteria were isolated from native soil taken from Shell Pond in 2016. Isolation methods are described in Xia et al. (2017). These strains were kept at 80°C in glycerol stocks. E.coli DH5α was used as a negative control. To revive each strain, a small amount of stock culture was incubated in 10 mL of LB at 37°C for three days. Each strain was plated on LB plates to create single colonies. From here, single colonies were selected and grown in 5 mL of LB for 16 hours.

### Nile Red stock solutions

1 mM and 0.1 mM stock solutions of Nile Red were made in DMSO. These stock solutions were encased in tin foil and kept in a −20°C freezer until use.

### Addition of Nile Red to crude oil

To determine whether concentration of Nile red had an effect on fluorescence intensity, we conducted several assays with three concentrations of Nile Red (0.1%, 0.5%, and 1%) for the 1 mM and 0.1 mM stock solutions. First, we sterilized the crude oil by autoclaving it three times and filtering it twice with 0.22 μm filters. Next, each concentration of Nile Red was incubated with 1 mL of crude oil for 20 minutes in 2 mL Eppendorf tubes. The Nile Red-crude oil solution was shaken periodically to assure the mixing of the two substances. We found that more consistent results were found with the 1% concentration of the 1 mM stock solution of Nile Red. This may be due to the low concentration of DMSO in the 1 mM stock solution, which allows for a more even complexing of the dye and the crude oil. We observed that the DMSO in the more diluted stock solution (0.1 mM) tended to pool at the bottom of the Eppendorf after complexing with the oil for 20 minutes. To determine the stability of the Nile Red-crude oil solutions, we left these solutions standing at room temperature in racks for 24 hours in full light. We observed a slight decrease in FI (roughly 20%), likely due to the fact that Nile Red is photo-sensitive and some bleaching may have occurred. As such, it is ideal to create new stock solutions of dyed crude oil before the start of each experiment.

### Assay set up

To set up each assay, 96 well plates were filled with crude oil which had been complexed with Nile Red as described above (ranging from 16 μl - 250 nl). For the control wells, LB or MSM was added to the wells until a total of 200 μl was reached. For wells containing bacteria, 100 μl of bacteria in LB or MSM at an OD of 0.6 were added to each well; LB or MSM was then added to bring the total volume of each well to 200 μl. Each treatment was replicated three times. Assays were sealed with parafilm and placed on a shaking incubator (120 rpm) in a dark room. To measure FI and OD, we used a Tecan plate reader. OD readings were taken at 600 nm in a circular pattern with 4×4 readings. FI readings were taken at excitation 535nm and emission 650 nm in a circular pattern with 6×6 readings. We measured FI and OD every day from T_0_-T_5_ and then again at T_10_ and T_20_.

### Staining of biosurfactant-based vesicles

Vesicles were stained with a 1 μg/mL stock solution of FM™ 4-64 Dye (*N*-(3-Triethylammoniumpropyl)-4-(6-(4-(Diethylamino) Phenyl) Hexatrienyl) Pyridinium Dibromide) which was made with a potassium phosphate buffer (pH 7). Briefly, 20 μl of culture was incubated with 20 μl of dye for 30 minutes. 1 μl of this solution was placed on a glass microscope slide and imaged with a fluorescent microscope.

### Microscopy

1ul aliquots of media from 96 well plates were placed on glass microscope slides and imaged using a Zeiss AxioImager M1 with a Hamamatsu Orca 03 12-bit grayscale digital color camera. Images were taken at 40x and 100x. Nile Red was detected using the Texas Red filter. To image intact biofilms within each well in glass-bottom 96-well microtiter plates, we used a Zeiss AxioObserver Z1 Live-Cell system with a QImaging Retiga SRV and a QImaging 5MPix Micropublisher camera. Images were taken with a 0.01-0.05 second exposure time (depending on magnification) and analyzed using the Zeiss Axio Vision software.

### Statistical analysis

We used t-tests to determine whether media type (LB or MSM) had a significant effect on crude oil fluorescence. We used one-way analysis of variance (ANOVA) to determine whether Nile Red concentration had a significant effect on crude oil fluorescence intensity. Differences among treatments were assessed by reference to the standard F tests. We also used ANOVA to determine whether microbial strain had an effect on degradation of crude oil. Crude oil depletion efficiency (%) was calculated by taking the FI at D_0_ and subtracting the FI at D_3_, divided by the FI at Day_0_. We used Pearson Correlation analysis to determine whether there was a correlation between FI and OD. To create a calibration curve to relate FI to amount of crude oil, we created linear models and compared these to quadratic and polynomial models of the data, followed by visual examination of the residuals of each model to evaluate the suitability of each model for explaining the data. We further tested the validity of the slope and y-intercept values for the model using t-tests. The confidence interval for each is β_o_ = b_o_ ± tsb_o_ and β_1_ = b_1_ ± tsb_1_ where sb_o_ and sb_1_ are the standard errors for the intercept and slope, respectively. To determine if there is a significant difference between the expected (β) and calculated (b) values we calculated t and compared it to its standard value for the correct number of degrees of freedom. We selected the model with the best ‘fit’ for the data based on R squared value and statistical significance. General statistics, ANOVA, and t-tests were conducted in R (v. 3.2.2, “Fire Safety”) using packages stats (v. 3.4, R core team) and psych (v. 1.6.4).

