## Supplementary material for "A high-throughput fluorescence-based assay for rapid identification of petroleum degrading bacteria": SI

K. E. French[1*] and N. Terry[1]

[1] Department of Plant and Microbial Biology, Koshland Hall, University of California Berkeley, Berkley, CA 94720

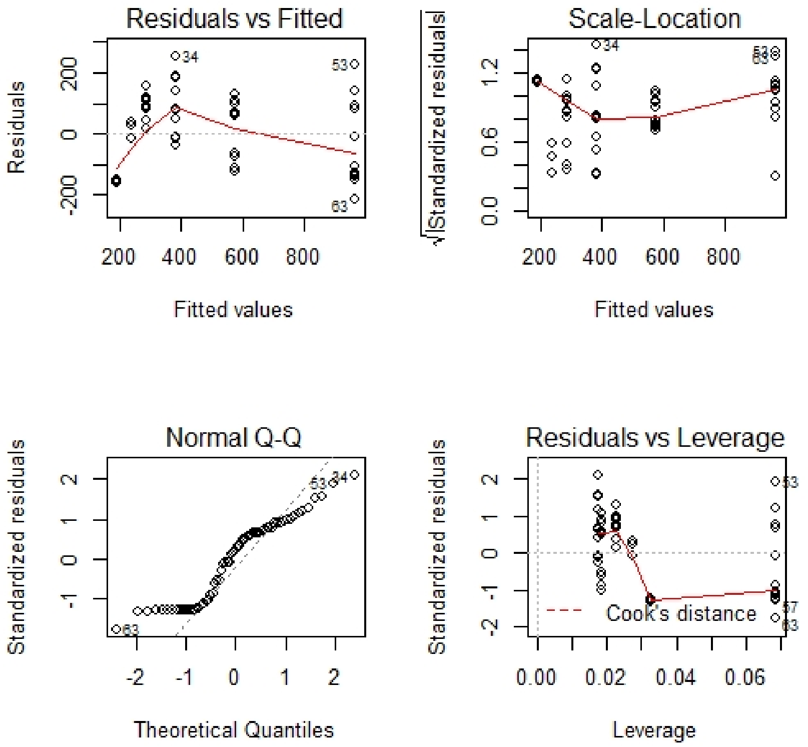


**Figure 1: Distribution of residual errors in calibration curve used to correlate fluorescence intensity with crude oil content.** The plot in the upper left shows the random distribution of the residual errors plotted versus their fitted values around a horizontal line representing a residual error of zero. The Q-Q plot in the lower left suggests the residual errors are normally distributed. The scale-location plot in the upper right shows the random distribution of the square root of the standardized residuals as a function of the fitted values. The plot in the lower right shows each point's leverage around contour lines for Cook's distance; all points are below distances of 0.5, indicating that removing a given observation has little effect on the regression results. Distances larger than 1 would indicate the presence of a possible outlier or a weak model. The analysis of residual and fitted errors corresponds to the data in Fig.1 C of the main text.

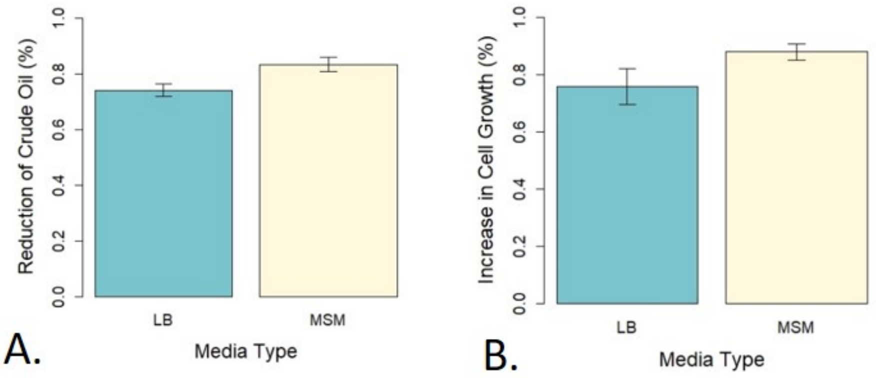


**Figure 2: Effect of media type (LB versus MSM) on bacterial degradation of crude oil (A) and cell growth (B).** Bacteria degraded more crude oil in the presence of MSM but there was no significant effect of media type on bacterial biomass (based on OD taken at 600nm).

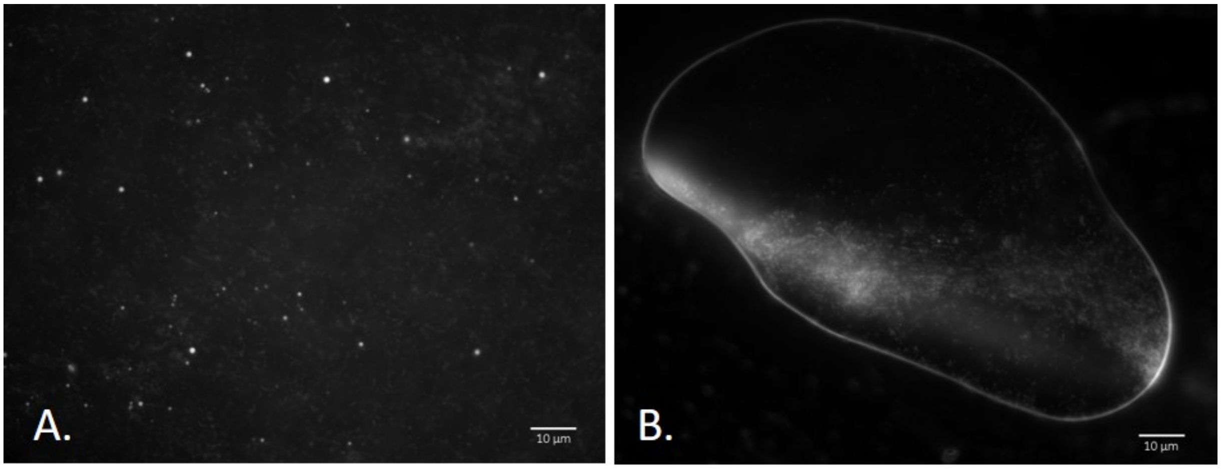


**Figure 3: Experimental evidence shows *Pseudomonas* sp. BSS produced membrane-bound vesicular structures containing crude oil.** To determine whether the spheres observed in earlier experiments were membrane bound (e.g. vesicles) we incubated cultures of *Pseudomonas* sp. BSS with un-dyed crude oil for 48 hours and then incubated the culture with the membrane dye FM 4-64. Bacterial membranes and vesicular structures were clearly marked by the membrane dye. The observed vesicular structures contained oil droplets (A) and sometimes bacteria and oil (B). The giant vesicle in B is 121.27 µm in diameter.

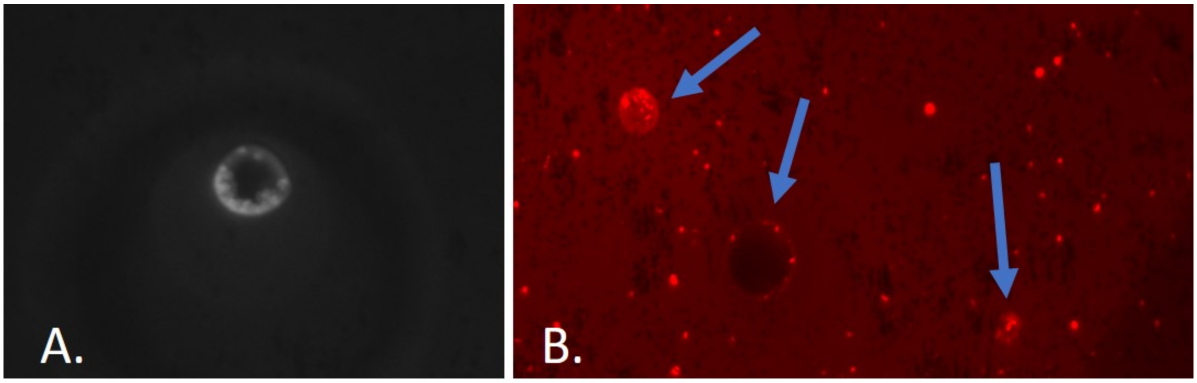


**Figure 4: Experimental evidence shows that other wild-type bacteria known to produce biosurfactants also produce vesicular structures when exposed to crude oil**. The wild-type consortia of bacteria known as FH 2-2 (isolated from Shell Pond, Bay Point, CA) produced vesicles ca. 10 µm wide when exposed to 2% crude oil (A, B). In the false color image (B), the arrows point to vesicles. The red dots and rods are bacteria. Bacteria were seen swimming inside the vesicles. Both images are derived from samples taken from the Nile Red assay. At $T_{10}$, a 20 µl aliquot was removed and incubated with 20 µl of membrane dye (FM 4-64); 1 µl of this solution was spotted onto a slide for microscopic analysis.

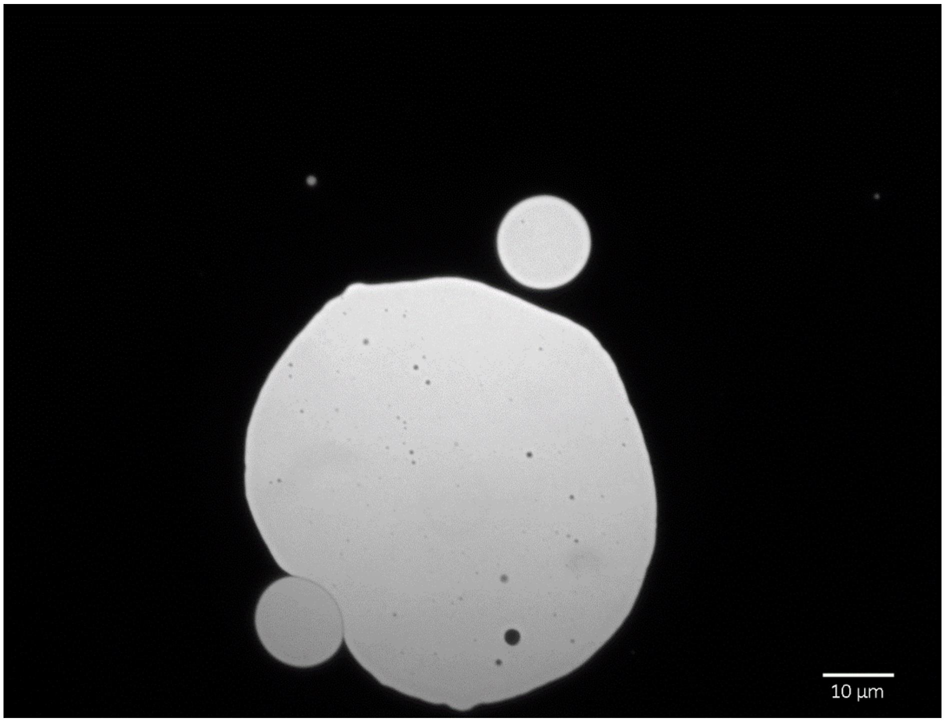


**Figure 5: Experimental evidence with the synthetic biosurfactant SDS shows that biosurfactants can spontaneously form vesicles when exposed to crude oil.** 50 µl of 1% SDS was added to 100 µl dH20 and exposed to 1 µl crude oil (not dyed with Nile Red). Smaller vesicles (5-10 µm in diameter) merged to form large vesicles >100 µm in diameter. Experiment details are described in the methods section.
